# Three-dimensional single-particle reconstruction by atomic force microscopy allows rapid structural-based validation of recombinant SARS-CoV-2 Spike protein from a single topology image

**DOI:** 10.1101/2025.05.20.655128

**Authors:** Liisa Lutter, David M. Beal, Maria Stanley, Joanne Roobol, Sarah Martin, James D. Budge, Phoebe E. Lee, Emi Nemoto-Smith, Ian Brown, Martin J. Warren, C. Mark Smales, Wei-Feng Xue

**Affiliations:** School of Natural Sciences, University of Kent, CT2 7NJ, Canterbury, UK; Quadram Institute Bioscience, Norwich Research Park, Norwich, NR4 7UQ, UK; National Institute for Bioprocessing Research and Training, Foster Avenue, Mount Merrion, Blackrock, Co. Dublin, A94 X099, Ireland

**Keywords:** atomic force microscopy, single-particle reconstruction, contact-point reconstruction, CPR-AFM, structural biology, cryo-electron microscopy, image analysis

## Abstract

Atomic force microscopy (AFM) is a versatile multi-modal imaging method frequently used for structural characterisation of biological surfaces at the nanoscale. However, AFM-based three-dimensional single-particle reconstruction has hitherto not been possible due to the tip-sample convolution artifact that distorts AFM images of individual molecules, and the disconnect between two-dimensional AFM images of surface deposited molecules and their three-dimensional structures. Here, three-dimensional single-particle analysis was developed for rapid structure-based validation of protein structures using as few as a single AFM topology image, based on contact-point reconstruction AFM (CPR-AFM) in an integrative approach with cryo-electron microscopy maps available in the Electron Microscopy Data Bank by template matching. This approach was demonstrated on the structural validation of recombinant trimeric ectodomain of SARS-CoV-2 Spike glycoprotein to show its immediate utility as a rapid structure-based sample quality control method in the recombinant expression and purification of the Spike protein samples that can be used in vaccines and therapeutics research. These results show that three-dimensional single-particle reconstruction by AFM is possible, that high signal-to-noise AFM imaging offers a rapid and cost-effective way of validation or identification of three-dimensional protein structures at single particle level, and that AFM can be linked to structural data derived from methods such as cryo-electron microscopy, resulting in integrative methodologies with new capabilities for structural biology.

## INTRODUCTION

Atomic force microscopy (AFM) is a scanning probe microscopy technique that can provide structural information of biomolecules deposited on a surface substrate by scanning of the sample surface with a physical nano-sized tip, thus mapping the 3D topography of the molecules of interest. It is a versatile multi-modal imaging method that can provide structural, mechanical, and chemical information of the scanned molecules, and is a well-established method for nanoscale characterisation for life-sciences and materials research (e.g. ^1–4^). AFM has recently embarked on a renaissance where contemporary developments have highlighted new unique capabilities that are particularly useful in structural biology studies of individual molecules ^5^. For example, contact-point reconstruction AFM (CPR-AFM) allows the three-dimensional (3D) tip-sample contact point clouds to be extracted from the two-dimensional (2D) topology images, thereby allowing 3D structural analysis of individually observed particles ^6,7^. Localisation AFM (L-AFM) allows the top contact points in molecular surfaces to be localised with near-atomic resolution based on high-speed AFM data ^8,9^. Simulation AFM (S-AFM) allows AFM images to be generated from structural data ^10,11^, models, or predictions ^12,13^, thereby facilitating comparison and template matching analyses and adding value to existing structural data in widely available databases and tools. Owing to the high signal-to-noise ratio of individual AFM topology images, the structures of individual molecules detected on the scanned images can be analysed, facilitating the use of AFM for rapid structure-based assessment of proteins on an individual-particle basis without the averaging requirement of other structural biology workflows such as in cryo-electron microscopy (cryo-EM) ^5^. Furthermore, the individual particle approach allows the structural heterogeneity of molecular populations to be assessed. This approach has been previously demonstrated on the three-dimensional (3D) reconstruction of individual self-assembled amyloid filaments ^6^, which subsequently enabled the quantification of the structural properties of individual filaments ^14^. Application of cryo-EM map-based S-AFM has also demonstrated how the complementary structural information from near-atomic resolution cryo-EM single-particle analysis data from ensemble- or subpopulation-averaged particles, and 3D reconstructions of individual filament envelopes from AFM data, can be integrated allowing new capability for structural identification of individual protein particles ^10,11^.

Despite these recent advances in AFM methodologies aimed at high-resolution structural biology applications, AFM imaging is not yet commonly used as a molecular structural biology tool because the tip-sample convolution artifact distorts AFM images of individual molecules. There is also a disconnect between two-dimensional AFM images of surface deposited molecules and their three-dimensional structures as molecules may not be observed from all angles by AFM. Thus, 3D single-particle reconstruction of protein molecules imaged by AFM, while conceptually similar compared to cryo-electron microscopy based single-particle reconstruction, has not hitherto been carried out. Here, we demonstrate that AFM-based 3D single-particle reconstruction is possible by combining experimental optimisations for the deposition of molecules onto the surface substrate, with 3D CPR-AFM based contact-point cloud analysis. This approach was demonstrated on the structural validation of recombinant trimeric ectodomain of SARS-CoV-2 Spike glycoprotein.

The severe acute respiratory syndrome coronavirus 2 (SARS-CoV-2) emerged as the pathogenic agent which led to the coronavirus disease (COVID-19) pandemic and global public health threat upon emergence in 2019 ^15,16^. The virus is characterised by a lipid envelope from which highly glycosylated Spike (S) transmembrane protein trimers protrude, forming a protein corona ^17^. Viral entry into host cells for enveloped viruses, including SARS-CoV-2, occurs through the fusion of the viral and host membranes, at the plasma membrane or via the endosomal pathway, which is energetically facilitated by conformation changes within viral fusion proteins ^18^. Host cell entry mechanisms for SARS-CoV-2 are primarily facilitated by the Spike (S) protein on the virion surfaces, which mediates host cell surface receptor angiotensin-converting enzyme 2 (ACE2) recognition and consequently viral entry into host cells ^19^.

Structural molecular biology of the S protein has been an important focus point for vaccines and therapeutics research. In this respect, molecular and structural biology workflows have enabled the rapid elucidation of structural information of key SARS-CoV-2 and human proteins involved in viral mechanisms of infection ^20^. In particular, cryo-EM single-particle reconstruction studies have revealed the trimeric structure of the Spike protein to high-resolution molecular detail (**Fig. 1**). Several hundred structural models of the Spike protein have now been deposited to the Protein Data Bank (PDB) and Electron Microscopy Data Bank (EMDB), revealing structural changes of the S protein with various mutations ^21^ and ligands ^22^ under different conditions. Cryo-EM studies have further revealed several prefusion conformational states of the S protein, in which up to two of the three receptor-binding domains (RBDs) can adopt an open conformation ^23^, as well as a structural continuum of the flexibility of RBD conformations ^24^. While cryo-EM single-particle reconstruction analyses have resulted in the acquisition of near-atomic resolution structural information from a wide variety of S protein samples, this level of structural detail is achieved through time- and resource-intensive facilities and workflows. In cases where the high-resolution structural information has been resolved and is widely accessible in the PDB and EMDB (e.g. for S protein), but where routine validation of the quality of folding of the protein for downstream biochemical or cellular studies is necessary, methods capable of providing rapid structure-based assessment of samples are important. This is of particular importance for COVID-19 research carried out with recombinantly expressed Spike protein ^25^ because, for example, the structure of prefusion-stabilised two-proline Spike (S2P), which has been widely used for both laboratory and clinical studies, degrades when stored at 4°C for longer than a week, leading to downstream impacts on the immunogenic response elicited in mice ^26^.

**Figure 1.**
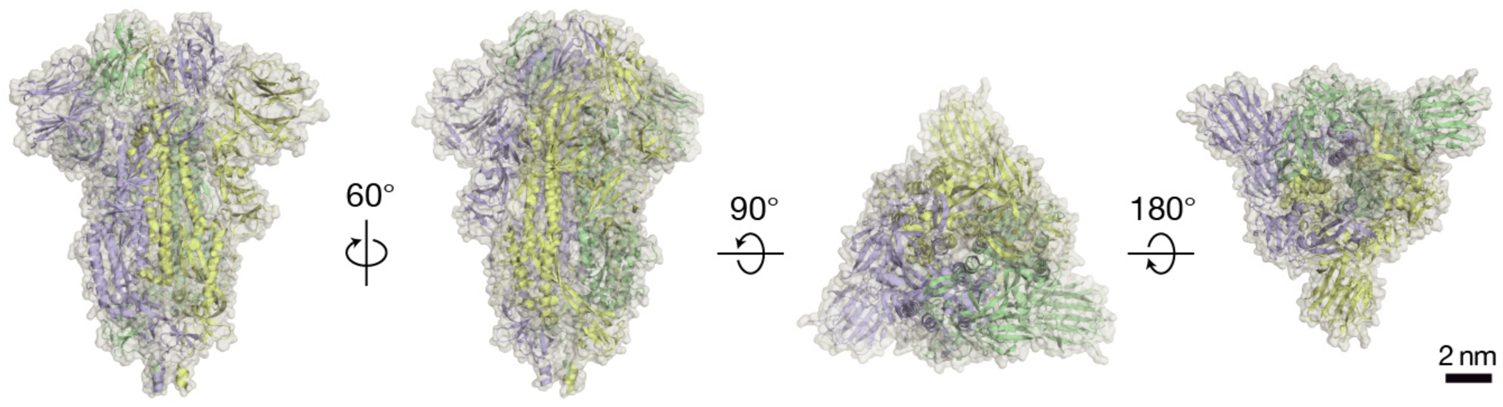
The trimeric ectodomain structure of the SARS-CoV-2 Spike protein has been resolved by cryo-EM to high-resolution molecular detail. Four different rotational views of a molecular model of S protein (PDB entry 6VXX ^20^) were visualised using PyMol. Each chain of the trimeric structure is coloured differently (yellow, green and purple, respectively). The scale bar represents 2 nm.

Here, we show that AFM height topology image data contain sufficient structural information for 3D single-particle analysis using images of individual Spike protein particles observed in a continuum of different orientations obtained by experimental optimization of molecular deposition conditions. A CPR-AFM analysis of the image data integrated with cryo-EM map-based template matching algorithm subsequently allowed the unknown orientational parameters for each extracted particle to be recovered. Subsequently, 3D single-particle reconstruction was then carried out that resulted in the reconstruction of the surface envelope of the S protein using as few as one single height topology AFM image. Thus, this approach facilitates rapid structure-based quality control for recombinantly expressed and purified proteins, and demonstrats its utility as a cost- and time-effective structure-based sample quality control method in the recombinant expression and purification of the S protein samples that can be used downstream in vaccines and therapeutics research.

## RESULTS

### Optimisation of molecular deposition conditions allows AFM topology imaging of individual S protein molecules in a continuum of different orientations

Nanoscale imaging of recombinantly expressed and purified ectodomain trimer S protein (expression and purification using established protocol described in the Methods section and **Supplementary Fig. S1**) was first deposited on fresh cleaved HOPG for AFM topology imaging. To facilitate subsequent 3D single-particle analysis, the deposition conditions were optimised so that individual S protein molecules can be observed in a wide range of orientations. Thus, the aim for the molecular deposition procedure was to partially embed individual S protein particles in a background buffer layer so that the individual molecules are resting in different orientations in a background raised from the HOPG surface floor. An analogy to this deposition strategy is to randomly toss small toys into a sandbox so that they are partially embedded in the sand but also partially exposed above the sand in different orientations. This approach to deposition was achieved by employing several experimental strategies. One, the molecules were deposited on a HOPG surface substrate with hydrophobic character, which aids in forming a background layer from the sample drying process. Two, undiluted stock sample solutions from the purification procedure were directly used in order to maximise sample concentration during the deposition procedure. Three, the deposition volume was small (10 μL) and was allowed to reach a semi-dry state over 10 min. Four, a small wash with 100 μL sterile-filtered MilliQ water was carried out prior to gentle drying with a stream of N_2_ gas to clean the surface layer of protein and buffer components so that some S protein molecules become partially exposed during the AFM topology scans. AFM imaging was subsequently carried out by gentle force-distance curve-based AFM imaging (see Methods section). The acquired AFM height topology images revealed a surface displaying uniformly distributed particles with triangular appearance in a variety of orientations (**Fig. 2a**).

**Figure 2.**
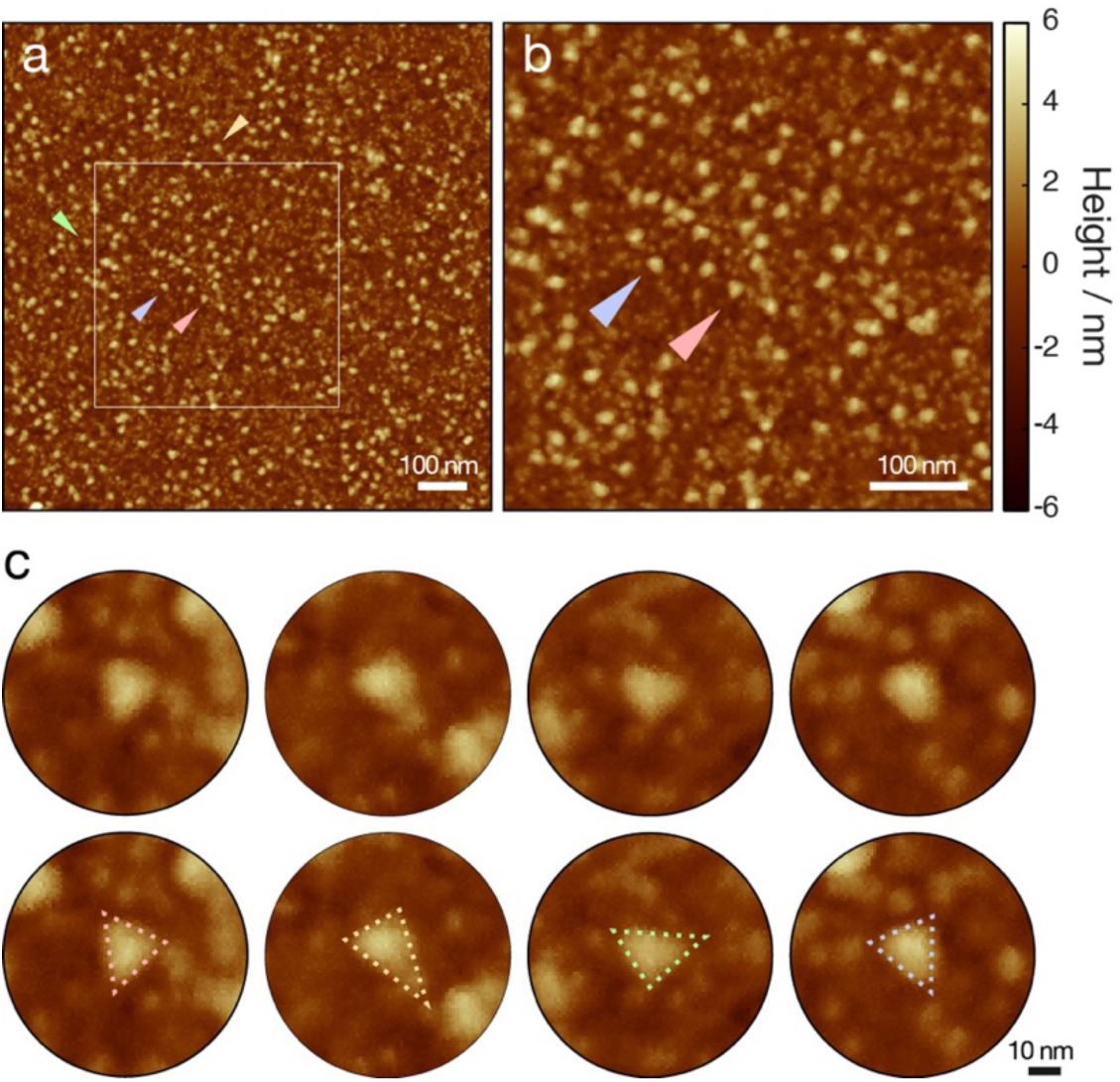
AFM imaging of SARS-CoV-2 S-protein particles deposited on HOPG results in height images with evenly distributed particles in different orientations. (a) Representative height topographic image of a 1024×1024 pixels, 1 μm × 1 μm area scan of the specimen surface. The scale bar indicates 100 nm. (b) A 2ξ magnified view of the area indicated by the white box in (a). The scale bar represents 100 nm. (c) Magnified views of example particles indicated by coloured arrowheads in the image from (a) and (b). The bottom row shows the same examples as the top row examples but with dashed lines indicating the triangular appearance of the particles. The scale bar represents 10 nm.

Next, individual particle images were picked and extracted from the AFM height images via an automated particle picking algorithm (**Supplementary Fig. S2**). A Laplacian of Gaussian convolution filter was applied to the AFM height images. Regional maxima were subsequently identified on the Laplacian of Gaussian filtered image and used as centre x- and y-coordinates to extract individual S protein particle images from the original unfiltered AFM image data. Since the apparent size of an AFM imaged S protein particle is expected to vary depending on the observed orientation of the particle, Laplacian of Gaussian kernels with different standard deviation values were tested. Optimum particle picking results, estimated from visual inspection, were obtained using a standard deviation value of 6 pixels. As an example, 582 particles were automatically picked on the image shown in **Fig. 2a**.

### Cryo-EM map-based S-AFM facilitates estimation of unknown orientational parameters of individual experimental particle images

Knowledge of the relative orientation of each individual particle observed on the AFM images is required in order to relate individual experimental particle images to each other in 3D space. Additionally, an estimate of the AFM probe tip radius is required to move recorded pixel x/y-coordinates and z-values of each pixel on the image to the tip-sample contact points in 3D spatial coordinates by CPR-AFM. Here, a template matching approach utilising existing cryo-EM based structural map of the trimeric ectodomain of S protein obtained from the EMDB (EMD-21452, ^20^) was used. A set of reference AFM images of the S protein were generated by a S-AFM method that simulates hard-sphere contacts between the probe tip and the iso-surface of the cryo-EM map using the contour level recommended by the authors of the map in the EMDB metadata (**Fig. 3**). The simulated reference image set contains simulated AFM images of the cryo-EM map rotated by a full 360° rotation along all three coordinate planes with a defined angular sampling frequency of 10° in every possible permutation. Simulations were carried out by identifying tip-sample contact points between the iso-surface of the rotated cryo-EM map and a model of the AFM probe tip with equivalent radius as the estimated tip radius of the probe used for experimental data collection. The simulated images also have identical x/y-sampling density compared with the experimental data.

**Figure 3.**
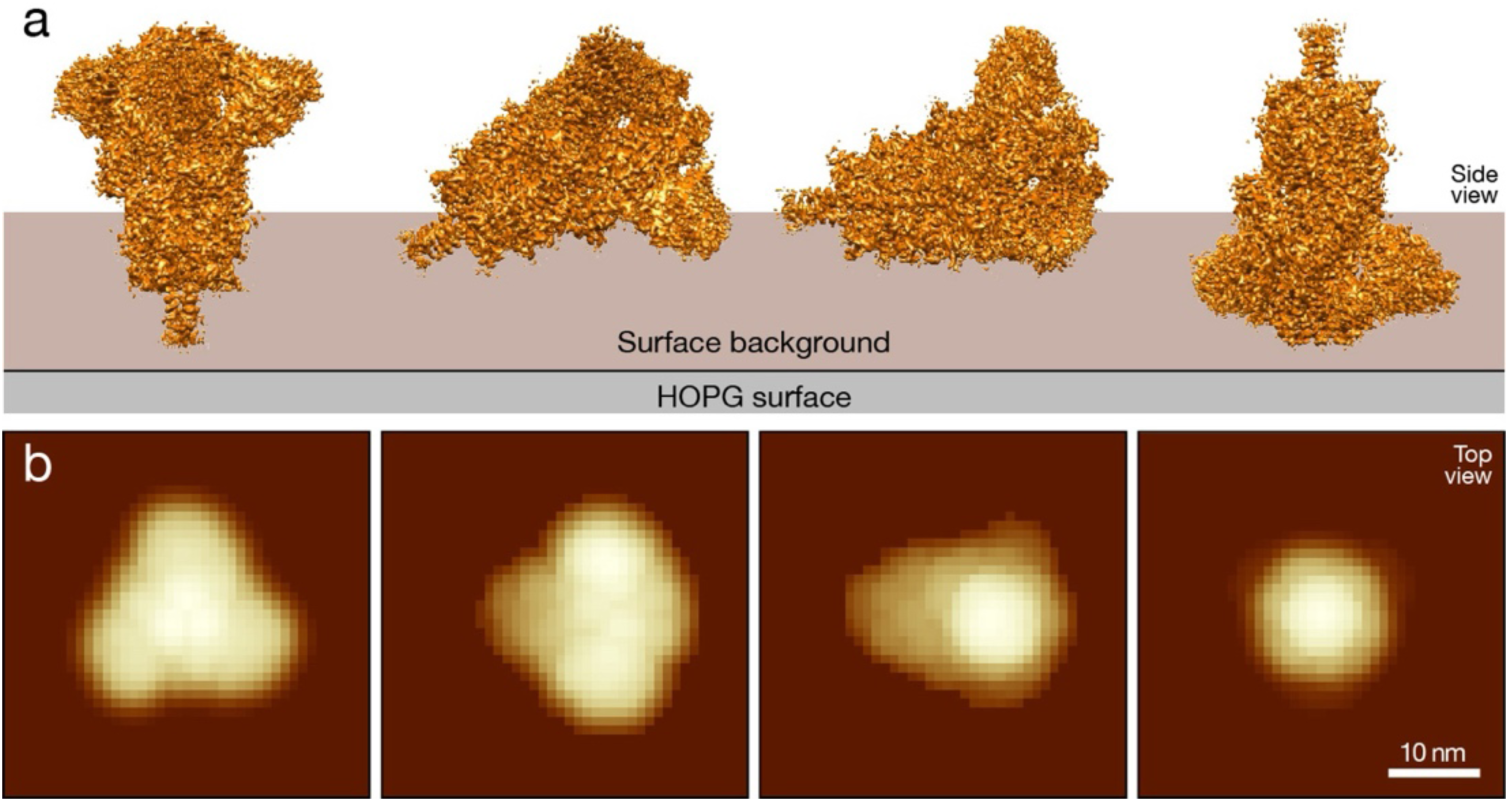
Cryo-EM map-based S-AFM allows generation of simulated reference image set of S protein in different orientations. (a) Schematic illustration of cryo-EM map iso-surface of the trimeric ectodomain of Spike protein (EMD-21452, ^20^) partially buried and resting on the sample specimen in four representative orientations. (b) Corresponding simulated AFM images of the S protein in the orientations shown above in (a) with the tip model and sampling pixel density equivalent to that of experimental AFM images. Scale bars represent 10 nm.

As the tip radius is initially unknown, an initial optimisation step is carried out in which normalised 2D cross-correlation values were calculated between the individual experimental images of the particles and a set of simulated reference images with varying tip radius. The tip radius was estimated to be the value that resulted in the highest cross-corelation value between the data and the simulated reference images. The AFM probe tip radius can significantly increase from their manufacturer-supplied nominal value as a result of the tip-sample contacts during the imaging process. For the image data of S protein particles, the estimated tip radius values were found to be higher than the nominal value of 2 nm given by the probe manufacturer in every image acquired. Four 1 μm × 1 μm AFM images with the smallest estimated tip radii (including the example image show in **Fig. 2a**) were selected for further analysis and their mean tip radius was estimated to be 3.4 nm. Images collected with larger tip radius led to an increased convolution effect and dilation of sample surface features, thus appearing more ‘blurry’ in comparison.

A full set of reference images generated from the cryo-EM map-based iso-surface of the trimeric S protein (EMD-21452 ^20^) and a 3.4 nm tip radius models were used for further analysis of the extracted AFM imaged Spike protein particles. These simulated topographic reference images of the S protein particles were used to estimate the rotational and translational parameters associated with the individual Spike protein particle images observed and extracted from the AFM image data through a templating matching strategy. Normalised 2D cross-correlation values were evaluated between each experimental particle image to every simulated reference image to test all possible orientational parameters. The simulated reference image with known orientational parameters that resulted in the highest cross-correlation value was identified for each experimental S protein particle image (**Fig. 4**). The orientational parameters of the highest scoring template reference image, and the x/y-alignment between this and the extracted experimental image allowed the orientational parameter of the individual experimental particle image, and subsequently its corresponding relative coordinates on the S protein in 3D space to be estimated. For every experimental particle image extracted from the AFM image data, the estimated orientations (rotational) and coordinate (translational) parameters from the template matching step were recorded.

**Figure 4:**
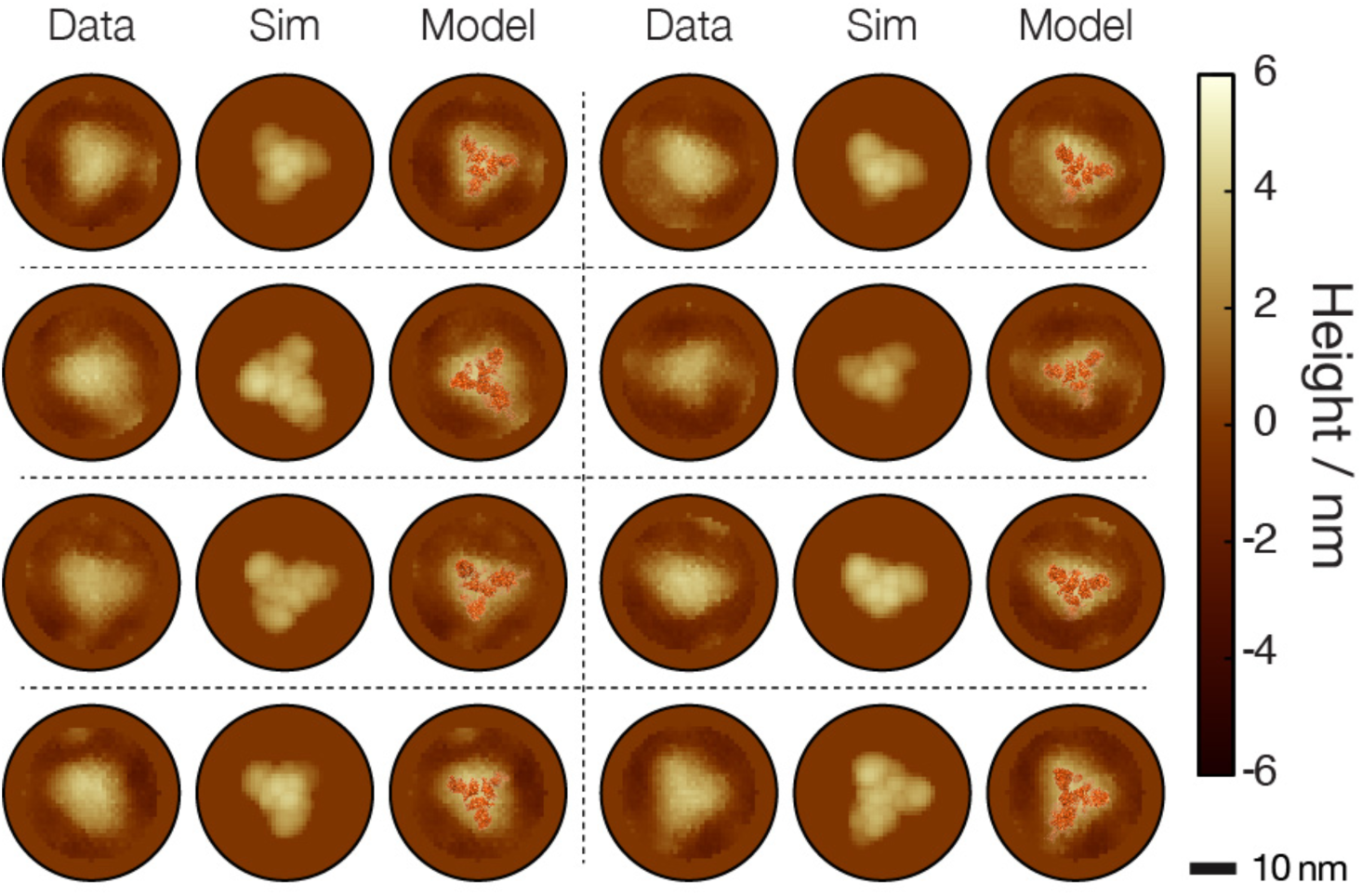
Estimation of particle orientations by template matching to simulated reference images. Template matching of the particle images extracted from the AFM data and simulated topology images of the rotational states based on cryo-EM map of S protein that best matched the data. Eight matched examples are displayed. For each example, the image data (left) are shown together with the best-match simulated reference image (centre) and a model of the S protein based on the cryo-EM map (EMD-21452 ^20^) in the best matched orientation superimposed onto the data (right). The scale bar represents 10 nm in all images.

In order to validate whether the quality of the experimental data and the template matching algorithm results in sufficient specific identification of the particles observed on the AFM images and their orientational states, two negative control reference data sets where generated and tested as templates. A trimeric HIV-1 Envelope protein (EMD-21111 ^27^) and a tetrameric plant nucleotide-binding leucine-rich repeat receptor RPP1 (EMD-30450 ^28^), both similar in dimensions to the trimeric ectodomain of S protein, were chosen to evaluate the specificity of the template matching procedure and the effect of similar template symmetry on the resulting cross-correlation scores. These negative control reference image sets were used as template in identical manner as the S protein reference images in the template matching procedure described above. Analysis of the RMSD between the extracted images of the particles seen on the experimental AFM data and the S protein reference images were 0.63 nm ± 0.01 nm (SE). For the negative control reference image sets, using the trimeric HIV-1 Envelope protein (EMD-21111) resulted in an RMSD of 0.77 nm ± 0.01 nm (SE) and using the tetrameric plant nucleotide-binding leucine-rich repeat receptor RPP1 (EMD-30450) resulted in and RMSD of 0.69 nm ± 0.01 nm (SE). These results showed that template matching between the experimental image data and the trimeric and tetrameric negative control reference data sets resulted in an overall increased RMSD values. Furthermore, the trimeric control data set did not lead to higher image matching scores, demonstrating that the template matching algorithm is not solely detecting the correct symmetry but is specific to the structural details of the reference data used as the template.

### 3D single particle reconstruction of the S protein surface envelope is possible from one single AFM topology image and facilitates rapid structure-based assessment

Finally, we carried out 3D single-particle reconstruction using the experimental particle images extracted from the AFM image data together with their individual estimated orientational and translational parameters. A total of 582 SARS-CoV-2 Spike protein particles were extracted from the example AFM images scanning a 1 × 1 μm area shown in **Fig. 2a**, and these particle images were initially tested for 3D single-particle reconstruction of the S protein surface envelope (**Fig. 5, Supplementary Movie S1**). The reconstruction algorithm operates by applying the orientational and translational operations estimated from the reference image that best matched each experimental AFM-imaged particle in opposite directions and in reverse order on the deconvoluted 3D contact-point cloud of each experimental particle image. Thus, for 3D single-particle reconstruction, 3D coordinates of the molecular surfaces in the form of a 3D contact-point cloud from each extracted and centred particle were first obtained using the CPR-AFM ^6,7^. The points in the point cloud higher than ¾ of the maximum height for each particle image were subsequently taken and translated on the x/y/z-axes in the opposite direction estimated from the translational parameters of its best-matched reference image. This was followed by rotation in the opposite direction and in the order of z-, y-, and x-axes using the rotational parameters estimated from its best-matched reference image. Analysis of the distribution of particle orientations (**Supplementary Fig. S3**) reveals a preference for side and top views, possibly leading to uneven local coverage of the surface envelope. Nevertheless, from particles extracted from one single topology height AFM image, a 3D contact-point cloud representing the surface envelope of the S protein was reconstructed. The contact-point cloud was subsequently visualised as a volumetric 3D density map, as well as 2D contact-point density image slices through its volumetric density map (**Fig. 5b-e, Supplementary Movie S1**).

**Figure 5.**
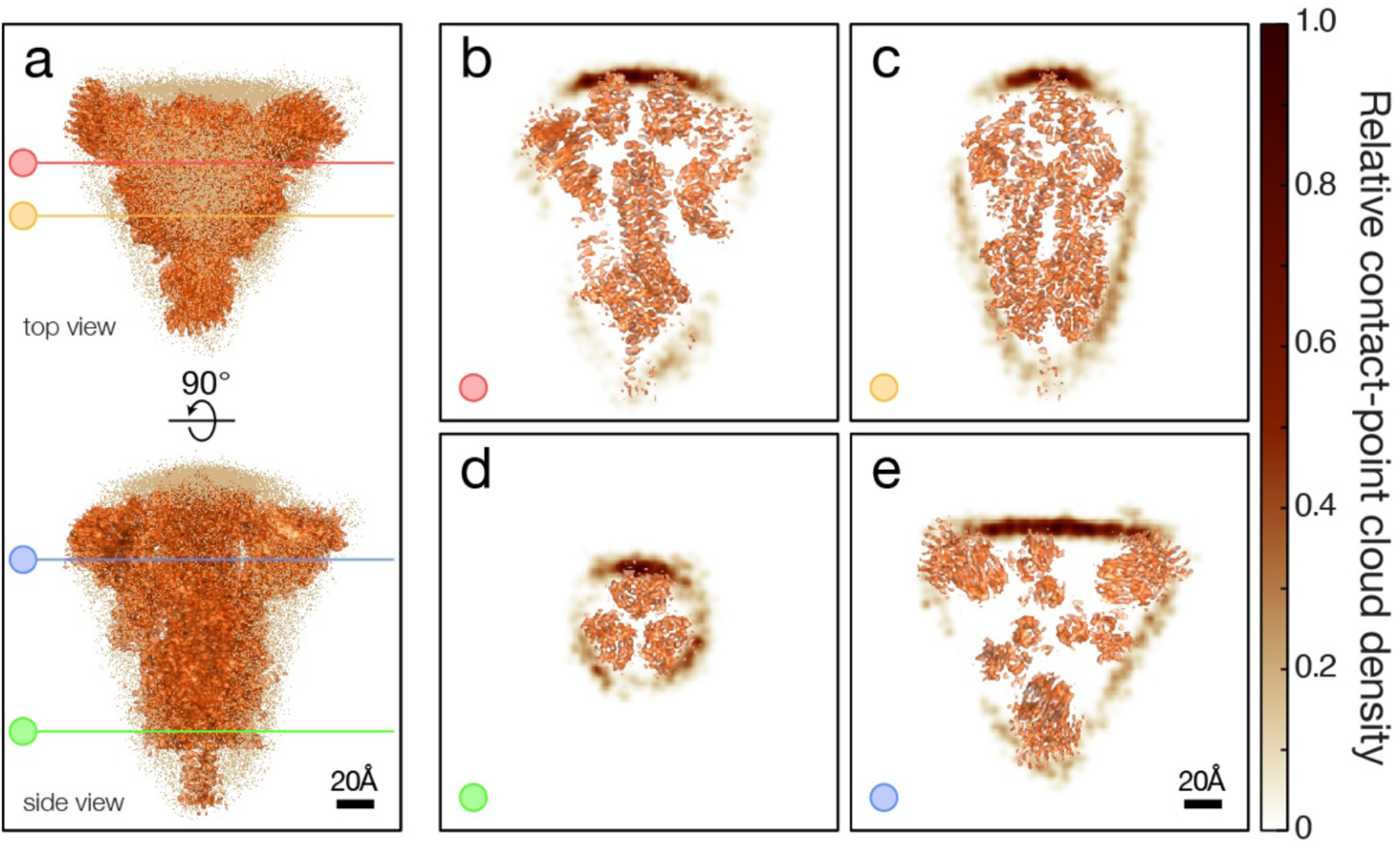
Single particle 3D reconstruction of a trimeric ectodomain of S protein from a single AFM height topology image. (a) 3D contact-point cloud (dots) reconstructed from 582 particles extracted from an experimental AFM image shown in Fig. 2a. Top and side views are shown together with cryo-EM structural map (EMD-21452 ^20^) of the trimeric ectodomain of S protein. Coloured lines indicate corresponding coordinates of contact-point density map slices shown in (b)-(e). The AFM-based surface envelope density maps are shown together with the cryo-EM map slices in **Supplementary Movie S1** and in (b)-(e) at locations indicated by the coloured slice lines in (a). The AFM-based densities along the sliced planes are represented by the colour intensity as shown in the colour bar. Scale bars represent 20Å in all images.

The 3D single-particle reconstruction of the WT SARS-CoV-2 S protein surface envelope from one single AFM height topology image showed that the molecular shape of the sample molecules closely corresponded to that of the Spike protein structure determined by cryo-EM (**Fig. 5**), establishing that AFM imaging-based rapid 3D single-particle reconstruction is possible. The resulting 3D contact-point map of the sample molecules exhibit the three-fold symmetry along the horizontal x/y-plane (**Fig. 5e, f**). The density slice planes further reveal the AFM based contact-point density closely followed that of the outer envelope of the S protein 3D map, with the protruding surface features exhibiting higher densities, as shown by the increased colour intensity. The slices show reduced or no AFM contact-point density in the centre of the S protein map, as expected, since these areas are inaccessible to contact with the AFM tip. The three outward projecting densities corresponding to the expected regions of the RBD domains (**Fig. 5a** top map) show variations in the intensity of the densities on the AFM based 3D reconstructed contact-point density map, which may be indicative of the presence of a mixture of open and closed RBD states. To validate the reproducibility of the single AFM topology image-based 3D single-particle reconstruction, 3D reconstruction was carried out independently using three other AFM images (with 761, 585 and 640 S protein particles picked, respectively) acquired with estimated average probe tip radius of 3.4 nm, comparable to that of the example image shown in **Fig. 2a**. The mean pairwise Chamfer distance (a pairwise similarity measure for point clouds) is 1.7Å for the four single AFM topology image-based 3D reconstructed contact-point clouds, confirming the reproducibility of the single-particle reconstruction workflow. These results provided a structure-based validation of the protein particles in the sample of recombinant S protein, and unequivocal demonstration that fast and resource-effective single AFM image-based 3D single particle reconstruction is possible.

## DISCUSSION

Atomic force microscopy topology imaging is a widely used imaging tool that allows images of excellent signal to noise ratio to be acquired. This is a feature that enables individual protein molecules to be imaged and seen even from a single observation on a single image. Here, we report the development of an algorithm to reconstruct the 3D protein surface envelope of a protein by 3D single-particle analysis using AFM height topology images. This algorithm is demonstrated on recombinantly expressed Spike protein particles from the original SARS-CoV-2 virions. The workflow allows the reconstruction of the 3D surface envelope from just one single AFM height image through a template matching approach using existing cryo-EM derived information. This integrative structural biology approach combines the rapid, accessible, and low-cost AFM topology imaging of individual molecules with the near atomic-resolution structural information provided by cryo-EM, allowing a rapid structure-based assessment of the sample molecules and add value to the wealth of data now publicly available in the EMDB. Such an integrative approach is also amenable to combinations with structural templates obtained from other structural biology methods, simulations, or artificial intelligence (AI) based structure prediction techniques. The shape and symmetry of the reconstructed 3D contact-point cloud closely corresponded to the shape and three-fold rotational symmetry of the folded trimeric S protein, and this is reproducible based on independent reconstructions from different individual AFM height topology images. Although there are 3D single-particle analyses reconstructed structural maps of the SARS-CoV-2 Spike using negative-stain electron microscopy ^29^, or to near-atomic resolution by cryo-electron microscopy, these approaches require data sets containing hundreds of thousands or more particle images. This report on AFM-based single-particle reconstruction of the S protein, on the other hand, shows that a rapid 3D single-particle analysis is achievable from just one single high signal-to-noise AFM height topology image. This also shows that sufficient structural information is available in the AFM height images and implies the feasibility of *ab initio* contact-point cloud reconstruction in future developments.

The simplicity and cost-effective approach to 3D single-particle reconstruction using AFM imaging we report here could be applied to rapid structure-based assessment of protein sample quality, for example for the S protein in the case of the development of recombinantly expressed protein-based vaccines, or protein based biotherapeutic drugs, where protein batch quality control and structural validation is of high importance. In the response to the COVID-19 pandemic, the importance of having a range of different vaccine technologies has become evident. Although mRNA vaccines have been widely used, a range of protein-based vaccine technology may be especially suitable for achieving vaccine equity by facilitating access in regions where maintaining storage and cold-chain conditions is challenging ^30^ or for overcoming vaccine hesitancy in regions with low uptake by providing options which may be perceived to be safer ^31^. Furthermore, the Spike protein is useful as a research tool for understanding the molecular basis of SARS-CoV-2 infection ^32,33^ but the stability and quality of the folded protein upon storage deteriorate with storage time^26^. Thus, validating the folded state of the protein samples used for downstream analyses is of fundamental importance. The AFM imaging and 3D structural reconstruction approach demonstrated here can provide such rapid information on the folding quality of the recombinantly expressed protein in a cost-effective manner.

The demonstrated 3D single-particle reconstruction algorithm for topographic AFM data establishes opportunities for further developments. For example, the integrative template matching approach linking to cryo-EM data can be extended to generate reference images from data sources other than cryo-EM, such as PDB molecular models from nuclear magnetic resonance spectroscopy (NMR) or X-ray crystallography studies. Furthermore, the recent advances in protein structure prediction (e.g. by AlphaFold ^34^) enables undertaking this integrated reconstruction approach on proteins for which no molecular model based on experimental data exists, through the incorporation of deep learning-based computational algorithms. In addition, multimodal 3D surface envelope reconstructions may be possible by mapping single-particle information on the mechanical, biophysical and chemical properties of the proteins from other AFM imaging modes (e.g. ^35–37^) or coupled techniques to the reconstructed 3D contact-point clouds. For example, nano-IR has been used in conjunction with AFM to determine surface properties of Spike protein particles at a nanometre spatial resolution ^38^. Finally, although the lateral resolution of AFM imaging and the tip-sample convolution effect limit the currently achievable 3D envelope resolution, further developments to AFM imaging and computational post-acquisition algorithms, for example by calculating detection probability from multiple scans of the same object in the same orientation by HS-AFM ^8^, will lead to improvements to the resolution of the 3D reconstruction of the contact-point cloud. This may make it possible to distinguish between different conformational states or protein-ligand complexes (**Fig. 6**), thus further facilitating the single-particle analysis of protein structures as well as structural analysis on an individual molecule level.

**Figure 6.**
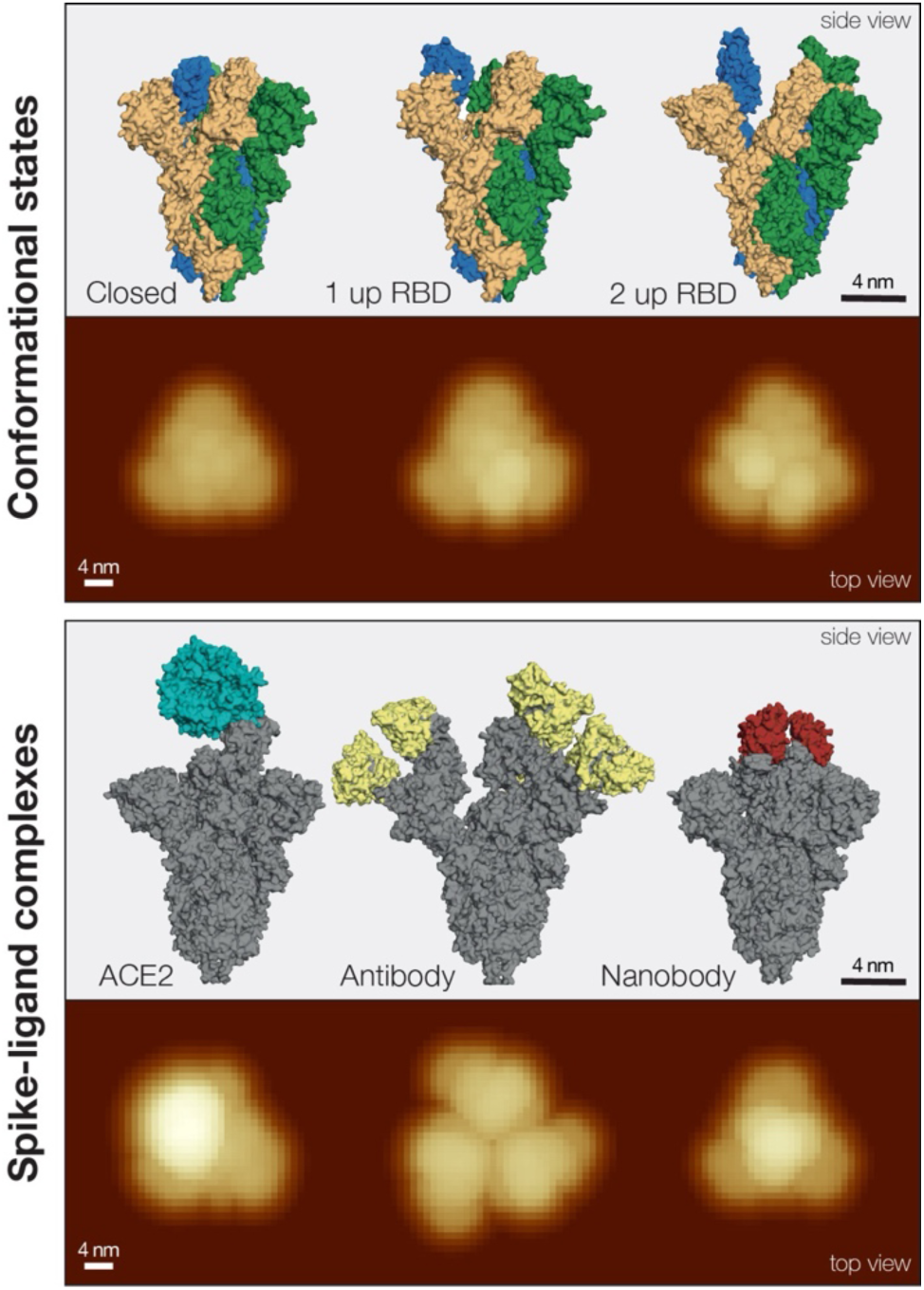
Conformational variations of the SARS-CoV-2 Spike protein and Spike protein-ligand complexes revealed by cryo-EM may be distinguished by template matching to simulated AFM images. The top box shows different conformational states of the S protein, from the closed state to one or two open receptor binding domains (RBDs). Each subunit of the trimeric protein is shown in a different colour. Simulated AFM topography images using the cryo-EM maps are shown below the structural models. The bottom box shows structural models of complexes formed by the Spike protein and various ligands, such as the ACE2 receptor, antibodies or nanobodies, respectively. The Spike protein trimer subunits are shown in dark grey, and the ligands are highlighted in blue, yellow or red for the three different ligands from left to right, respectively. Simulated AFM topography images based on the corresponding cryo-EM maps are shown bellow the structural models. All structural models were visualised using PyMol from the PDB entries 7KDG ^39^, 6VYB ^20^, and 7A93 ^40^ (conformational states; left to right, corresponding to EMDB entries EMD-22821, EMD-21457, EMD-11683, respectively), and 7DF4 ^23^, 7CWU ^41^, and 7OAN ^42^ (Spike-ligand complexes; left to right, corresponding to EMDB entries EMD-30661, EMD-30488 and EMD-12777, respectively). The S-AFM images were simulated using comparable tip radius (3.4 nm) and identical pixel density as the AFM image data (Fig. 2). The scale bars represent 4 nm in all panels.

## METHODS

### Cloning of the Spike protein-encoding gene and transfection and expression of Spike protein in CHO-S cells

The gene sequence for the SARS-CoV-2 Spike (S) protein was that kindly provided by the Wrapp laboratory ^43^. For this study, the sequence was commercially synthesised (GeneART Invitrogen, USA) after being codon optimised for expression in Chinese hamster ovary (CHO) cells such that it coded for the same protein sequence as described by Wrapp *et al* with the native ER signal peptide, removal of the transmembrane domain to allow secretion of material out of the cell, introduction of proline residues to aid stability and removal of cleavage sites ^43^. A C-terminal His-tag was added to the sequence for purification and detection purposes. The gene sequence was then cloned into the multiple cloning site of the commercially available pcDNA3.1 hygro plasmid (ThermoFisher).

Spike protein expression was undertaken in CHO-S cells (Gibco, Catalog number A1155701), maintained in CD-CHO media (Gibco) supplemented with 8 mM L-glutamine. CHO-S cells were transiently transfected with plasmid DNA encoding the full-length spike protein (CHO optimised) as follows: 100 µL plasmid DNA (at a concentration of 400 ng/µL in TE buffer) was gently mixed with 1×10^7^ CHO-S cells in 700 µL of CD-CHO media in a 4 mm gap width Gene Pulser® cuvette (Bio-Rad 165-2088), placed in a GenePulser Xcell electroporator and then electroporated as previously described ^25^. The cells were then transferred to 20 mL of CD CHO media. This process was typically repeated 15 times to give a final culture volume of 300 mL in a 1000 mL Erlenmeyer vented cap flask. The culture was then placed at 37°C in a shaking incubator (140 rpm) at 5% CO_2_ and left for 6 days.

### Recombinant Spike protein purification

Six days post-transfection, the secreted Spike protein was harvested by centrifugation at 1000×*g* for 10 minutes. The supernatant was then dialysed using dialysis cellulose membrane with a MW cut-off of ∼14000 Da (Sigma D9402-100FT) at a 1:10 ratio in 20 mM Tris-HCl pH 8, 500 mM NaCl for 2 × 2 hours at 4℃ with constant stirring (buffer was exchanged for fresh buffer after 2 hours). Chelating Sepharose Fast Flow (Cytiva 17057502) was loaded into an empty column, allowed to settle and then washed with 50 mL of dd H_2_O by gravity flow. Then, 50 mL of 50 mM NiSO_4_ was passed through the column. The column was equilibrated with 50 mL binding buffer (20 mM Tris-HCl pH 8, 500 mM NaCl, 5 mM imidazole). Following dialysis, 12 mL of Chelating Sepharose Fast Flow pre-charged with NiSO_4_ was added to the supernatant and stirred slowly at 4°C for 30 mins. The supernatant/resin mix was loaded gradually into an empty column and allowed to flow through as the resin settled. Once all the supernatant was loaded, the resin was washed with 100 mL of binding buffer, followed by 50 mL of 20 mM Tris-HCl pH 8, 500 mM NaCl, 30 mM imidazole and then 50 mL of 20 mM Tris-HCl pH 8, 500 mM NaCl, 50 mM imidazole. The His-tagged protein was eluted from the column with 50 mL of 20 mM Tris-HCl pH8, 500 mM NaCl, 400 mM imidazole. The 50 mL elution was concentrated to 1 mL using an Amicon™ Ultra-15 Centrifugal Filter Unit (Merck UFC910024) with a 100 kDa MW cut-off. The concentrated protein was then buffer exchanged into 2 mL 10 mM phosphate buffer pH 8, 100 mM NaCl using a PD10 de-salting column (Cytiva 17085101). Purified protein was stored at 4°C.

### Western blot analysis of purified Spike protein

Purified Spike protein samples were analysed by western blot using a mouse anti-polyhistidine primary antibody (Sigma H1029) and an anti-mouse alkaline phosphatase conjugated secondary antibody (Promega S3721) and developed using SIGMAFAST™ BCIP®/NBT substrate (Sigma B5655).

### AFM imaging and image processing

A 10 μl drop of the purified Spike protein sample was deposited onto freshly cleaved HOPG surfaces substrate (Agar scientific, AGG3389) and incubated for 20 minutes. Following the incubation, the semi-dry HOPG surface was washed with 100 μL of filter sterilised milli-Q water and then dried using a gentle stream of nitrogen gas. The sample specimens were imaged using a Multimode 8 AFM with a Nanoscope V controller (Bruker) operating under peak-force tapping mode in air using ScanAsyst probes (silicon nitride triangular cantilevers with tip height = 2.5-2.8 μm, nominal tip radius = 2 nm, nominal spring constant 0.4 N/m, Bruker). Each collected image had a scan size of 1 x 1 μm and 1024 × 1024 pixels. Nanoscope analysis software (Version 1.5, Bruker) were used to process the image data by 2nd order flattening of the height topology data to remove tilt and scanner bow.

Automated peak detection was carried out by initially applying a Laplacian of Gaussian filter in order to achieve local smoothing of the topographic data and facilitate peak picking. Each image was filtered independently using Laplacian of Gaussian kernels with standard deviation of 6 pixels. Subsequently, local maxima were found from the Laplacian of Gaussian filtered images by identifying pixels surrounded only by pixels with a lower value but are higher than a threshold value of 95% of the maximal value of the image matrix. The x- and y-coordinates of these local maxima were then recorded and used as centre points for extracting particles on the original non-filtered images.

### Simulation AFM of reference topographic images from cryo-EM maps

Cryo-EM density map of the Spike protein were downloaded from the Electron Microscopy Data Bank (EMDB). The volumetric Coulomb density map data (EMD-21452 ^20^) were imported into Matlab where all subsequent processing and analysis was carried out. The iso-surfaces of the cryo-EM density maps were found by applying the contour value provided by the authors of each of the EMDB data entries. For simulation of reference images, a model of the AFM tip with matching the estimated tip radius used in the imaging experiments was constructed and used to find the contact points between the tip model and the iso-surface coordinates when the tip is positioned at the x/y pixel-grid constructed with the pixel density equivalent to that of the experimental AFM data. As the tip radius on each experimental AFM image was initially unknown, it was estimated by an initial optimisation algorithm, which maximised the 2D normalised cross-correlation between extracted experimental particles and simulated images, while varying the radius of the tip model used for simulations. For tip radius optimisation, reference images were simulated by 60° sampling of iso-surface rotation on the three coordinate planes. For this, the iso-surface was systematically rotated by the x/y/z rotational angles followed by translations along the x/y-axes to centre it on the origin, before proceeding to identify contact points between the tip model and the iso-surface vertices. The three rotational and translational parameters corresponding to each simulated reference image were recorded. The optimised tip radius was then used to generate a tip model for subsequent simulation of a more extensively sampled set of reference images from the cryo-EM iso-surface, with 10° sampling on each of the three rotational axes. This reference data set was then used as the template in a template matching procedure with experimental particle images for 3D reconstruction of the S protein envelope. Simulated reference image data sets were also generated from two other cryo-EM maps (EMD-21111 ^27^ and EMD-30450 ^28^) using identical procedure, and subsequently used in control experiments to test the specificity of the template image matching algorithm.

### Topographic template image matching and single-particle 3D surface envelope reconstruction

The unknown orientation information of the input particle images was recovered through topographic template matching with a reference image set consisting of simulated images with known rotational and translational parameters. The normalised 2D cross-correlation between each input particle and each simulated image was calculated with varying x- and y-axis offsets, to account for non-centred input particles. For each input image, the rotational and translational parameters corresponding to the reference image that results in the highest normalised 2D cross-correlation between the input particle image data and the reference image are then recorded. The x- and y-axis offsets that result in the maximal correlation value are used to centre the input particle image.

Reconstruction of the 3D surface envelope from experimental AFM images was subsequently carried out by first estimating the tip-sample contact points in 3D space for each extracted particle image. These extracted x/y/z-coordinates for each particle were then translated in the opposite direction by the three translational parameters identified via template matching with simulated images of known rotational and translational parameters. Translation of the 3D coordinates was followed by rotation using the three rotational parameters also identified during the template matching procedure, in the opposite direction and in reverse order. The resulting reconstructed 3D point cloud was then visualised by constructing a contact-point density map using Gaussian kernel density estimation method with an average scalar bandwidth obtained with a non-parametric method ^44^ times the identity matrix. Slice planes orthogonal to the x/y-y/z- and x/z-axes of the volumetric density map of the reconstructed 3D point cloud were also visualised.

## Supporting information

Lutter et al Supplementary Movie S1

## ACKNOWLEDGEMENTS

This work was part funded via the Biotechnology and Biological Sciences Research Council (BBSRC) via grant BB/V011324/1 (CMS and MJW) and BB/Z516880/1 (WFX), Wellcome Trust Henry Wellcome Fellowship grant (209171/Z/17/Z) (DMB), and Engineering and Physical Sciences Research Council (EPSRC), UK DTP grant EP/R513246/1 (LL).

## SUPPLEMENTARY FIGURES

**Supplementary Figure S1.**
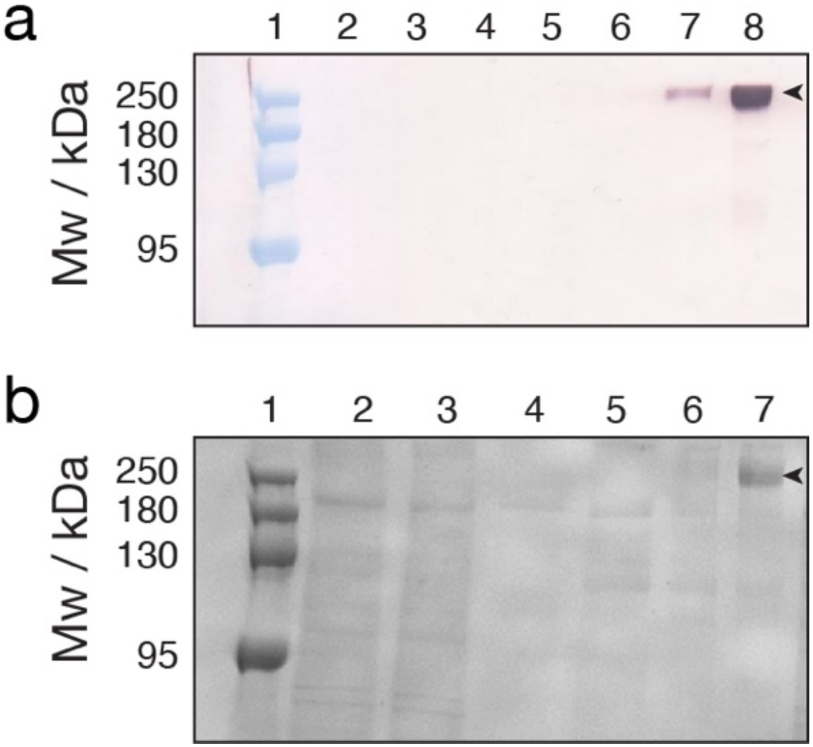
Western blot and SDS-PAGE analysis of the purified recombinant Spike protein produced and secreted from CHO-S cells. (a) Western blot analysis of the purified Spike protein. Lanes contain: 1) Molecular weight marker; 2) cell culture supernatant after dialysis; 3) flowthrough after loading column; 4) binding buffer flowthrough; 5) flowthrough from 30 mM imidazole wash buffer; 6) flowthrough from 50 mM imidazole wash buffer; 7) eluted protein in 50 mL elution buffer; 8) concentrated and buffer exchanged eluted protein. (b) Coomassie stained SDS-PAGE analysis of the purified Spike protein produced and secreted from CHO-S cells. Lanes contain: 1) Molecular weight marker; 2) cell culture supernatant after dialysis; 3) flowthrough after loading column; 4) binding buffer flowthrough; 5) flowthrough from 30 mM imidazole wash buffer; 6) flowthrough from 50 mM imidazole wash buffer; 7) concentrated and buffer exchanged eluted protein. Spike protein bands are indicated with an arrow.

**Supplementary Figure S2.**
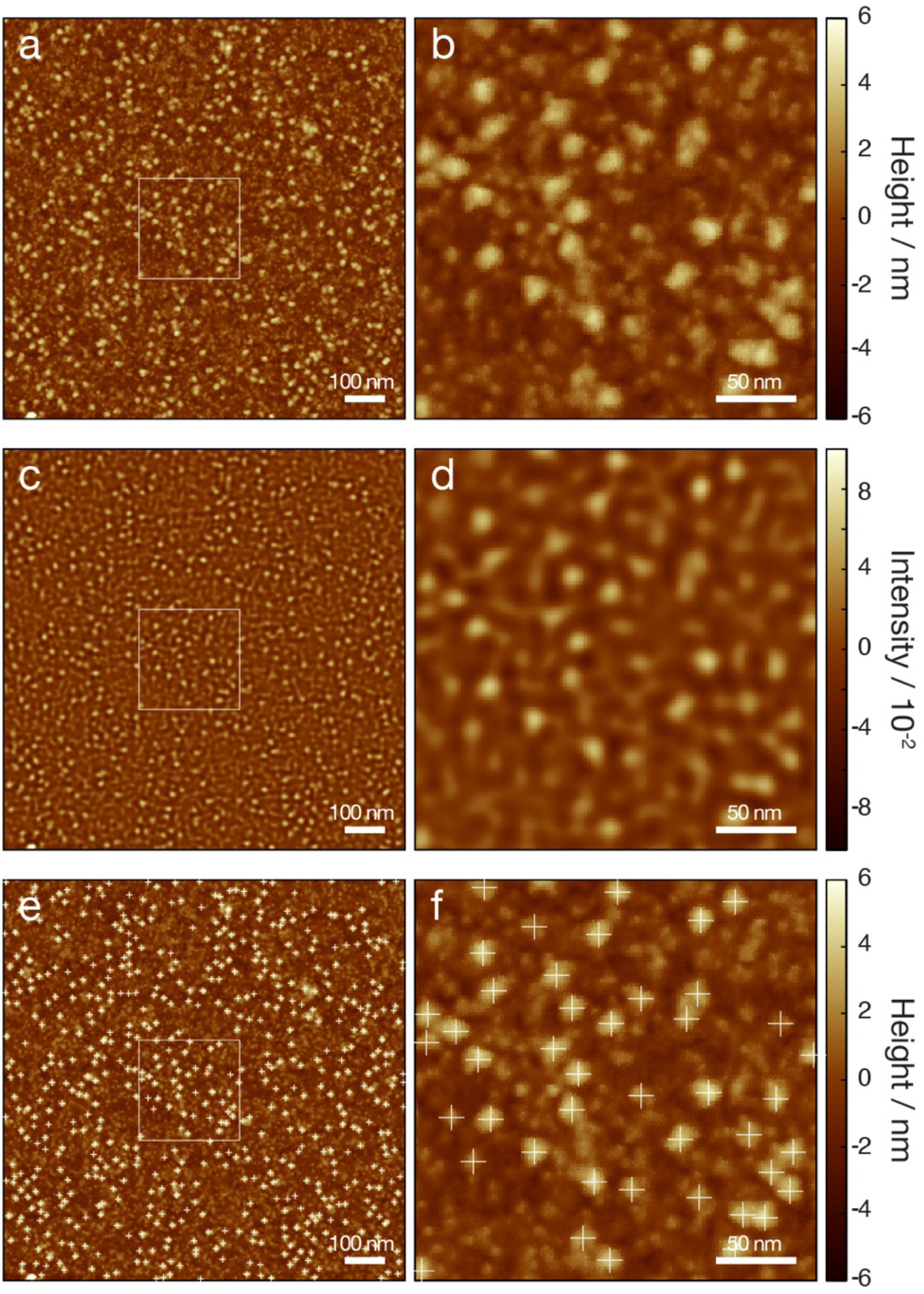
Automated and objective picking of S protein particle images by a Laplacian of Gaussian convolution image filter. (a) Representative height topographic image of a 1024×1024 pixels, 1 μm × 1 μm area scan of the specimen surface. The image is the same as shown in Fig. 2a. (b) A 4× magnified view of the area indicated by the white box in (a). (c) Laplacian of Gaussian filtered image of (a). (d) A 4× magnified view of the area indicated by the white box in (c). (e) The same height topology image as (a) with the x- and y-coordinates of detected particles labelled with crosses. These coordinates are subsequently used for extraction of the particle images. (f) A 4× magnified view of the area indicated by the white box in (e). The scale bars represent 100 nm in (a), (c) and (e), and 50 nm in (b), (d) and (f).

**Supplementary Figure S3.**
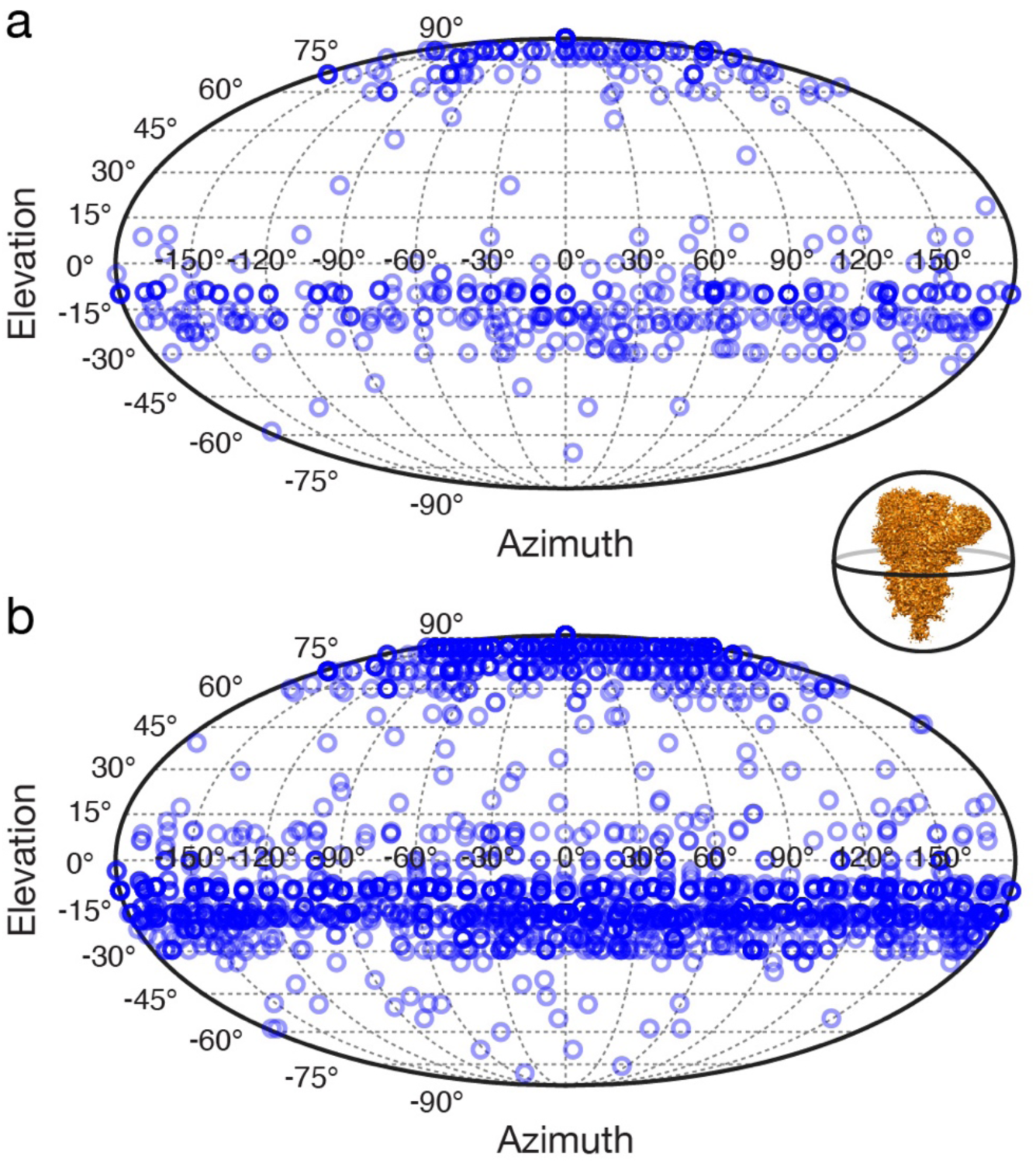
Distributions of orientations for particles picked and used in the 3D single-particle surface envelope reconstruction. (a) Azimuth and elevation angles for 582 particles picked from image shown in Fig. 2a. (b) Azimuth and elevation angles for all 2568 particles picked from all images. Inset shows the reference orientation for the Spike protein, which is the original orientation of the EMDB map EMD-21452 of the S protein ^20^. The Azimuth and elevation angles for individual particle image are show as circles projected onto an equal-area Mollweide projection of a spherical surface.

## Notes

### Competing Interest Statement

The authors have declared no competing interest.

